# IRON DEFICIENCY RESHAPES THE POPLAR ROOT AND EXUDATE METABOLITE PROFILES AND DRIVES EXUDATE-MEDIATED IRON MOBILIZATION

**DOI:** 10.64898/2026.10.16.759646

**Authors:** Maria Teresa Cieschi, Nadine Bouzari, Sylvie Citerne, Ana Alvarez-Fernandez, David Touboul, Sébastien Thomine

## Abstract

Iron (Fe) deficiency under alkaline conditions limits Fe bioavailability and reshapes plant metabolism. We asked how Fe deficiency at pH 7.5 alters the metabolite profiles of roots and root exudates of *Populus* × *canescens*, and whether exudates mobilize Fe at high pH. Hydroponically grown plants were deprived of Fe for 20 and 27 days. Roots and exudates, collected in nutrient solution and in distilled water, were profiled by UHPLC–QTOF MS, and metabolites were annotated using MS/MS molecular networking (MetGem) and public spectral libraries. Fe deficiency caused significant global shifts in both root and exudate metabolomes. In roots, phenylpropanoids increased at 27 days (proanthocyanidins, neochlorogenic acid, naringenin), whereas the salicylate glycoside salicortin decreased, suggesting a redirection of phenylpropanoid flux. In nutrient-solution exudates, eight flavonoids, including quercitrin, accumulated strongly at 27 days, and catecholic coumarins such as fraxetin were detected in roots and exudates. Exudates from Fe-deficient plants increased Fe mobilization from Fe(III) oxides at pH 7.5, and quercitrin solubilized Fe as efficiently as fraxetin. Alkaline Fe deficiency thus reconfigures poplar root and exudate metabolomes, and exuded flavonoids, not only coumarins, may contribute to Fe mobilization at high pH.

**HIGHLIGHT:** Iron deficiency at alkaline pH reshapes poplar root and exudate metabolomes; exuded flavonoids such as quercitrin, alongside coumarins, contribute to iron mobilization from insoluble iron oxides.

## INTRODUCTION

Iron (Fe) is indispensable for photosynthesis, respiration, and numerous redox reactions (Liang, 2022). At alkaline pH, however, Fe bioavailability declines sharply due to the precipitation as poorly soluble Fe (III) (oxyhydr)oxides. To cope with low Fe availability in the environment, land plants deploy two main strategies to acquire Fe: Strategy I—used by most dicots—combines rhizosphere acidification, Fe (III) reduction and Fe (II) uptake, whereas Strategy II—used by grasses—involves the release of mugineic acid family phytosiderophores into the rhizosphere, which form Fe (III) chelates that are subsequently taken up by root cells (Grillet and Schmidt, 2019). In calcareous environments, Strategy I species often complement their Fe-acquisition machinery by releasing Fe-mobilizing metabolites into the rhizosphere (Liang, 2022).

Among such metabolites, catecholic coumarins have emerged as key actors in Fe acquisition at high pH (Liang, 2022). In *Arabidopsis*, the phenylpropanoid pathway yields fraxetin via scopoletin 8-hydroxylase (Siwinska et al., 2018; Tsai et al., 2018) and produces the downstream redox-active metabolite sideretin (Rajniak et al., 2018). These coumarins mobilize Fe by chelation and/or reduction (Rajniak et al., 2018; Tsai et al., 2018), and fraxetin remains effective at alkaline pH where reductive chemistry wanes (Tsai et al., 2018), and recent evidence indicates that Fe (III)–fraxetin complexes can be directly taken up by roots independently of IRT1 (Robe et al., 2025). Coumarin secretion is tuned by external pH and supports Fe nutrition (Tsai et al., 2018; Paffrath et al., 2024). Root-secreted coumarins also modulate rhizosphere microbiota under Fe limitation (Harbort et al., 2020).

Beyond coumarins, catechol-bearing flavonoids—such as quercetin, luteolin, and catechins—can chelate Fe through catechol and other oxygen-containing coordination sites (Malacaria et al., 2021). O-methylation or other modifications of hydroxyl groups involved in Fe coordination generally reduce chelating and redox capacity, whereas free catechol groups favor Fe binding. Although their chemistry suggests a potential role in Fe mobilization at high pH (Kumar et al., 2024; Walencik et al., 2024), the contribution of root-released flavonoids to this process remains unclear.

Poplar (*Populus spp.*) is a fast-growing tree cultivated in short-rotation systems that supply wood for pulp and paper, veneers/engineered wood, and increasingly bioenergy/biorefinery feedstocks—placing it at the forestry–bioeconomy interface (FAO-IPC, 2024). Poplar also holds a model-tree status: the first tree genome, *Populus trichocarpa*, catalyzed tree genomics and mechanistic studies on wood formation and plant–microbe interactions (Tuskan et al., 2006; Jansson and Douglas, 2007).

Root exudates—including organic acids, phenolics, flavonoids, lipids, peptides, and N-containing metabolites—collectively orchestrate rhizosphere chemistry and shape its microbiota (Afridi et al., 2023). Root exudates serve as a primary carbon and energy source for microorganisms in the rhizosphere. In a study of a poplar bioenergy plantation, root exudates specifically contributed 22% of the estimated below-ground carbon inputs. However, the percentage of carbon in poplar root exudates can vary depending on environmental conditions and plant stress (Berhongaray et al., 2019). This interaction helps shape the root-associated microbiome and can enhance nutrient acquisition for plants (Camli-Saunders and Villouta, 2025; de Haas et al., 2025). Since plants grown in hydroponic systems often exude more total carbon than those in soil, the metabolites present in the nutrient solution would provide abundant substrates for microbial communities (Heuermann et al., 2023). For example, two poplar species (*Populus tremuloides* Michx. and *P. trichocarpa* Torr. & A.Gray ex Hook) exposed hydroponically to aluminum exuded organic acids, which accounted for 20–64% of total released in root exudates (Naik et al., 2009).

The functional relevance of root exudates is also increasingly documented in poplar, including their interactions with rhizosphere microbial communities (M. Li et al., 2022). However, how Fe deficiency at alkaline pH reshapes root metabolism and exudate composition remains largely unexplored in woody species.

Here, we use grey poplar (*Populus × canescens*), a natural hybrid of *P. alba* and *P. tremula*, as a woody experimental system to investigate how Fe deficiency at pH 7.5 alters the metabolomic profiles of roots and root exudates. We reveal the presence of catecholic coumarins in root exudates and increased accumulation of different classes of flavonoids, including flavones and flavonols, in nutrient-solution exudates under Fe deficiency. Beyond metabolite profiling, we evaluate whether poplar root exudates and selected exuded metabolites can mobilize Fe from Fe (III) oxides at alkaline pH.

## 2 MATERIALS AND METHODS

### 2.1 Reagents

All of the reagents used were of analytical grade, and solutions were prepared with type I grade water according to ISO 3696:1987, free of organic contaminants (Millipore, Milford, MA, USA).

### 2.2 Plant material and growth conditions

*In vitro* multiplication: Stem cuttings (1–2 cm long) from ca. 8-week-old plantlets of the poplar INRA 717-1-B4 clone, *Populus × canescens* (*P. tremula × P. alba*), were transferred to glass tubes containing 0.01 L of Murashige and Skoog (M&S) medium (Murashige and Skoog, 1962), whose composition is described in Supplementary Table S1. The glass tubes were covered with a metal cap and incubated in a growth chamber under controlled environmental conditions (21–18 °C, 63% relative air humidity, 16h/8h light/dark cycle). After 4 weeks, the plantlets had reached a height of approximately 4–6 cm and developed 4-5 leaves. Six plantlets were transferred to 2.0 L Araponic boxes (http://www.araponics.com, Liège, Belgium) containing Hoagland nutrient solution (NS) with the following composition: macronutrients (in mM), 1.50 Ca (NO_3_)_2_, 1.25 KNO_3_, 0.75 MgSO_4_, and 0.28 KH_2_PO_4_; micronutrients (in μM), 10.0 FeHBED, 5.00 MnSO_4_, 0.50 CuSO_4_, 1.00 ZnSO_4_, 50.0 KCl, 25.0 H_3_BO_3_, and 0.10 Na_2_MoO_4_; 3.00 Mm HEPES. The pH was adjusted to 7.5 with KOH 1.00 M. During the 2-week acclimation period, plantlets were covered by a glass vessel, and the NS was renewed weekly. Three weeks after transfer to the NS, Fe deficiency treatments were initiated. Plants were transferred in groups to beakers containing 400 ml of NS. Control plants received the NS described above, whereas Fe-deficient plants received the same solution without Fe (III)-HBED. Beakers were covered with black plastic containers to avoid light exposure of the roots. Each treatment consisted of five biological replicates (beakers) with three plants per replicate (n=5 beakers), and replicates were randomized across positions within the growth chamber. Two independent experiments were conducted, with plants exposed to Fe deficiency for 20 or 27 days. Leaf chlorophyll status was monitored every 2-3 days after the onset of Fe deficiency using a SPAD-502 chlorophyll meter (Minolta, Osaka, Japan). The pH of the NS was measured with a pH meter every day during Fe-deficiency period and before NS renewal.

### 2.3 Plant sampling and micronutrient analysis

Roots, stems, and leaves were separated and weighed after harvest and washed with ultrapure water. A portion of each tissue was ground with liquid nitrogen in a ceramic mortar and stored at -80 °C until analysis. The remaining material was dried in a forced-air oven at 60 °C for 2 days. Dried samples were milled, and 0.05 g of material digested using 2ml of HNO_3_ (65%, Sigma Aldrich) and 1ml of H_2_O_2_ (30%, Sigma Aldrich) for 6h at 120°C. Iron, manganese and zinc concentrations were determined by microwave plasma atomic emission spectroscopy (MP-AES; 4200 MP-AE, Agilent Technologies).

### 2.4 Hormone analysis of roots

Frozen root powder (3 mg) was extracted with 0.8 mL of acetone/water/acetic acid (80/19/1, %v/v/v). Stable isotope-labelled abscisic acid (ABA), salicylic acid (SA), jasmonic acid (JA) and indole-3-acetic acid (IAA) used as internal standards, were prepared as described in (Le Roux et al., 2014). One ng of each internal standard was added to the sample. The extract was vigorously shaken for 1 min, sonicated for 1 min at 25 Hz, shaken for 10 min at 10°C in a Thermomixer (Eppendorf®), and then centrifuged (8,000 x g, 10 °C, 10 min). The supernatants were collected, and the pellets were re-extracted twice with 0.4 mL of the same extraction solution, followed by vigorous shaking (1 min), sonication (1 min, 25 Hz), and centrifugation as described above. The three supernatants were pooled and dried.

Each dry extract was reconstituted in 100 µL of acetonitrile/water (50/50, v/v), filtered, and analyzed by ultra-high-performance liquid chromatograph coupled to electrospray ionization tandem mass spectrometry (UHPLC-ESI-MS/MS) using an Acquity UPLC system coupled to a Xevo TQ-S triple quadrupole mass spectrometer (Waters Corporation, Milford, MA, USA). Compounds were separated on a reverse-phase column (Uptisphere C18 UP3HDO, 100 x 2.1 mm, 3µm particle size; Interchim, France) using a flow rate of 0.4 mL min^-1^. The mobile phase consisted of (A) 0.1 % (v/v) acetic acid in water and (B) 0.1 % (v/v) acetic acid in acetonitrile. The column temperature was maintained at 40 °C. The following gradient was used (time, % A): 0 min, 98 %; 3 min, 70 %; 7.5 min, 50 %; 8.5 min, 5 %; 9.6 min, 0%; 13.2 min, 98 %; and 15.7 min, 98 %.

Mass spectrometric detection was performed in multiple reaction monitoring mode, using positive ionization for IAA and in negative ionization for ABA, SA and JA. The capillary voltage was set to 1.5 kV in negative ion mode, and the source and desolvation temperatures were 130 °C and 500 °C, respectively. Nitrogen was used as the cone and desolvation gas at flow rates of 150 and 800 L h^-1^, respectively, and argon was used as the collision gas at a flow of 0.18 mL min^-1^.

### 2.5 Root exudate collection and processing

To evaluate the effect of Fe deficiency on the metabolites released by poplar roots, root exudates in nutrient solution (RENS) were collected for each treatment daily during the last seven days of the experiment, filtered through filter paper, pooled in glass bottles, and stored at -20°C. For each batch, matrix blanks were obtained following the same procedure from identical beakers containing nutrient solution but no plants, maintained in parallel under the same experimental conditions and following the same collection procedure. Samples were then concentrated using Sep-Pack C18 cartridges (Waters, Mildford, MA, USA), eluted with 1 ml of 100% methanol, and stored at -80°C until analysis. At 20 and 27 days of Fe deficiency, roots were washed with pure water and transferred to beakers containing 400 mL of sterile ultrapure water. After, 12 h, root exudates in water (REDW) were filtered through filter paper, transferred to 500-mL glass spherical beakers, and frozen at -80°C. Matrix blanks (REDW from beakers without plants) accompanied each batch. Samples were freeze-dried and resuspended with 3.0 mL of ultrapure water. To separate low-molecular-weight compounds from mucilage, samples were centrifuged through 3-kDa Amicon-Ultra-4 mL centrifugal tubes (Merck, Darmstadt, Germany) during 30 min at 11,000 rpm and 4 °C using an Eppendorf Centrifuge 5804 R (Eppendorf, Hamburg, Germany).

### 2.6 Untargeted metabolomic analysis of roots and root exudates

Frozen root powder (ca. 50 mg) was extracted three times with 0.6 ml of cold 100% methanol by shaking in a Retsch MM400 mill (Restch, Düsseldorf, Germany) at 30Hz for 30 s. After each extraction, samples were centrifugated at 13,000 rpm for 3min at 4°C, and the supernatants were collected and pooled. The combined extracts were vacuum-dried in a Savant Speed Vac Concentrator (Thermo Fisher Scientific, Waltham, Massachusetts, MA, USA) and reconstituted in 500 µl of water containing 10% acetonitrile and 0.1% formic acid. Samples were then diluted to a final concentration of 1 µg ml^-1^ and filtered through 0.45 µm PVDF Ultrafree-MC centrifugal filter units (Millipore) prior to untargeted metabolite profiling.

RENS and REDW extracts, prepared and stored as described above, were thawed, vortex-mixed and aliquots of 300 and 300 µL, respectively, were vacuum-dried using a Savant Speed Vac Concentrator (Thermo Fisher Scientific). The residues were reconstituted in 500 µl of water containing 10% acetonitrile and 0.1% formic acid. Samples were then diluted to a final concentration of 1 µg ml^-1^ and filtered through 0.45 µm PVDF Ultrafree-MC centrifugal filter units (Millipore) prior to untargeted metabolite profiling.

The analysis was performed on a 1260 Infinity II UHPLC system (Agilent Technologies, Waldbronn, Germany) coupled to a quadrupole time-of-flight (QTOF) mass spectrometer (6546 Q-TOF, Agilent Technologies). Chromatographic separation was performed on an SB-C18-Poroshell column (2.1 mm × 150 mm, 2.7 µm, Agilent Technologies) with a mobile phase consisting of H_2_O/formic acid (99.9/0.1, v/v) (A) and acetonitrile (B). The column oven was set at 45°C. Compounds were eluted at a flow rate of 0.4 mL min^−1^ using a gradient from 5% to 100% B over 20 min, followed by 100% B for 3 min. The injection volume was 5 µL for all analyses. Mass spectra were acquired in positive ion mode. The electrospray source was operated under the following conditions: gas temperature, 325°C; drying gas flow rate, 10 L min^−1^; nebulizer pressure, 30 psi; sheath gas temperature, 350°C; sheath gas flow rate, 10 L min^−1^; capillary voltage, 3,500 V; nozzle voltage, 500 V; fragmentor voltage, 130 V; skimmer voltage, 45 V; and Octopole 1 RF voltage, 750 V. The use of a calibration solution, containing two internal reference masses (purine, C_5_H_4_N_4_, *m/z* 121.0509, and HP-921 [hexakis-(1H,1H,3H-tetrafluoropentoxy) phosphazene], C_18_H_18_O_6_N_3_P_3_F_24_, *m/z* 922.0098), routinely resulted in mass accuracy below 3 ppm. Data-dependent MS/MS events were acquired for the five most intense ions detected by full-scan MS over the *m/z* 200– 1,200 range, using an absolute threshold of 1,000 counts. Selected precursor ions were fragmented with a fixed collision energy of 20 eV and with an isolation window of 1.3 amu. The mass range of the precursor and fragment ions was set to *m/z* 200–1,200 and 50-1,200, respectively.

### 2.7 Untargeted metabolomic data processing and molecular networking

MSconvert software (Chambers et al., 2012) belonging to Proteowizard package (https://proteowizard.sourceforge.io/tools/msconvert.html, accessed on 19 October 2024) was used for data conversion from Agilent proprietary format to mzML files. Data preprocessing was performed with MZMine3 software (Schmid et al., 2023) with the following parameters: mass detection noise level was set to 1000 for MS and 50 for MS/MS; chromatogram building was achieved with the automated data analysis pipeline (ADAP) chromatogram builder algorithm, with a minimum of 3 scans per group, a minimum intensity of 10,000 and *m/z* tolerance of 0.005 (or 15 ppm). Chromatogram deconvolution was achieved with the ADAP wavelets algorithm with the following parameters: signal/noise threshold 30, minimum feature height 10,000, coefficient threshold 10, peak duration range between 0.00 and 1, RT wavelet range from 0.02 to 0.09, *m/z* center calculated as the median, *m/z* range for MS/MS of 0.1 and RT range for MS/MS of 0.2. Feature lists were deisotoped with the isotopic peak’s grouper algorithm with an *m/z* tolerance of 0.005 or 15 ppm, and an RT tolerance of 0.1 min. Peak alignment was performed using the join aligner method with *m/z* tolerance of 0.005 or 15 ppm, *m/z* weight of 80% and an RT tolerance of 0.1 min with a weight of 20%. Aligned chromatograms metadata were used for statistical analyses, first with all MS features. Then, MS/MS data were filtered, and chromatograms and metadata with only MS/MS data were exported to GNPS-FBMN compatible MGF and CSV files. Files were processed with MetGem 1.5.2 software (Olivon et al., 2018) with the following parameters: *m/z* tolerance was set at 0.04 and a minimum of two matched peaks was set for similarity between two nodes. All peaks in a +/−17 Da window around the *m/z* precursor were removed. Only peaks above 1% of the highest or the six highest peaks in a +/−17 Da window throughout the spectrum were kept. t-SNE visualization was constructed with the following parameters: at least one cosine score above 0.70 was requested. Number of iterations, perplexity, learning rate and early exaggeration were set to 10,000, 6, 10 and 12, respectively. To cope with the high number of MS/MS features, Barnes–Hut approximation angle was used and set at 0.7. Annotations were performed with GNPS (https://gnps.ucsd.edu/ProteoSAFe/libraries.jsp, accessed on 19 October 2024) and MS-DIAL (http://prime.psc.riken.jp/compms/msdial/main.html#MSP, accessed on 19 October 2024) public standards databases. For library searches in the GNPS (Schmid et al., 2021), MS-Dial (Takeda et al., 2024) and ISDB (Allard et al., 2016) databases, parameters were as follows: *m/z* tolerance = 0.2 Th; minimum matched peaks = 2; minimum intensity = 0%; *m/z* parent tolerance = 17 Th; min. cosine score = 0.60.

### 2.8 Complementary LC-MS analysis for metabolite annotation

Frozen root powder (ca. 100 mg) was extracted three times with 1 ml of cold 70% LC-MS grade methanol by shaking in a Retsch MM400 mill (Restch, Düsseldorf, Germany) at 30Hz for 5 min. After each extraction, samples were centrifugated at 13,000 rpm for 3 min at 4°C, and the supernatants were collected and pooled. The combined extracts were vacuum-dried in a Speed Vac Concentrator (MiVac, GeneVacTM, Ipswich, UK), the residue immediately reconstituted in 250 µL of water containing 5% methanol and 0.1% formic acid., and then filtered through 0.22 µm PVDF Ultrafree-MC centrifugal filter units (Millipore) prior to LC-MS analysis.

REDW extracts, prepared and stored as described above, were thawed, vortex-mixed and 300 µL aliquots were vacuum-dried using the miVac concentrator. The residues were reconstituted in 500 µl of water containing 5% methanol and 0.1% formic acid, filtered through 0.22 µm PVDF Ultrafree-MC centrifugal filter units (Millipore), and subjected to LC-MS analysis.

Samples were analyzed using an UHPLC (ELUTE, Bruker Daltonics, Bremen, Germany) coupled to a quadrupole time-of-flight mass spectrometer (compact QTOF, Bruker Daltonics) equipped with an electrospray ionization (ESI) source. Metabolites were separated on a Bruker Intensity Solo C18 column (2 µm, 2.1 × 100 mm) maintained at 30 °C. The mobile phases consisted of 0.1% formic acid in water (A) and 0.1% formic acid in methanol (B), delivered at a flow rate of 0.5 mL min⁻¹ using the following gradient: 5% B at 0 min, increased to 90% B at 15 min, and held at 90% B until 19 min. Samples were maintained at 4 °C in the autosampler, and 5 µL was injected for each analysis. The ESI source was operated with a capillary voltage of 3500 V, nebulizer gas pressure of 3 bar, dry gas (N₂) flow of 9 L min⁻¹, and dry gas temperature of 200 °C. Mass spectra were acquired in positive mode ion mode over an *m/z* range of 50– 1000. Selected precursors ions were fragmented with a collision energy of 35 eV and with an isolation window of 0.5 Da. Instrument control and data acquisition were performed using Bruker Compass HyStar 5.1 (version 5.18.1) and otofControl (version 5.2) software (Bruker Daltonics). For this analysis, root and REDW samples from control and Fe-deficient plants were pooled across biological replicates and the two sampling times (20 and 27 days) and analysed once. Coumarins were identified by comparing retention time, accurate mass and MS/MS spectra with those of authentic standards from an in-house library (fraxetin, fraxin, isofraxidin, fraxinol and scopoletin).

### 2.9 *In vitro* solubilization of Fe (III)-oxide by root exudates and individual phenolics

The Fe (III)-oxide solubilization assay was performed as in Sisó-Terraza et al. (2016), with minor modifications. Aliquots of 500 µL of REDW and RENS extracts from Fe-deficient plants, were dried for 2 h using the previously described SpeedVac concentrator, and the residues resuspended in 300 µL of 5 mM HEPES-KOH buffer (pH 7.5). For the assay, 10 mg of poorly crystalline Fe (III)-oxide, prepared according to Pérez-Sanz and Lucena (1995), or ferrihydrite prepared as described by Cieschi et al. (2021), were incubated with 1.5 mL of assay solution for 6 h in the dark at 25°C with continuous shaking (300 rpm) using a Thermomixer Comfort (Eppendorf AG, Hamburg, Germany). The assay solution contained either 100 µM of the tested phenolic compound or 300 µL of the root exudate samples, together with 600 µM ferrozine (as Fe (II)-trapping agent), 500 µM β-nicotinamide adenine dinucleotide phosphate (NADPH, serving as the electron donor), and was buffered with 5 mM HEPES-KOH (pH 7.5).

Following incubation, samples were centrifuged at 10,000 x g for 1 min, and the supernatants were filtered through 0.22 µm PVDF membranes. The filtrates were analyzed for Fe (II) content by measuring ferrozine absorbance at 562 nm. Total Fe concentrations were subsequently determined by atomic emission spectrometry using an MP AES 4200 system (Agilent, USA). Prior to analysis, 900 µL of each filtrate was mineralized by addition of 100 µL ultrapure 65% HNO₃ (TraceSELECT Ultra, Sigma–Aldrich).

### 2.10 Statistical data analysis

Metal content data were analyzed separately for each tissue (leaves, roots) and sampling time (20 and 27 days). Hormone content data were analyzed for roots and sampling time (20 and 27 days). Outliers were identified by the iterative Grubbs test (α = 0.05). Normality and homoscedasticity were assessed by the Shapiro-Wilk and Levene tests, respectively. Differences between Control and Fe-Def were evaluated by Student’s t-test when both assumptions were met, or by the Mann-Whitney U test otherwise, with significance set at *p* < 0.05. All analyses were performed in Python (v3.11) using the SciPy library (v1.11). The processed feature table (CSV) was imported into MetaboAnalyst 6.0 (Pang et al., 2024; www.metaboanalyst.ca, accessed 19 October 2024). Features were normalized by sample median, log10-transformed, and mean-centered (“mean scaling”). Univariate comparisons between Fe-deficient and control groups were performed per feature using two-sided Student’s *t*-tests with α = 0.05. Fold change was computed as log2 (Fe-deficient / Control) from the non-log-transformed intensities (i.e., pre-log values); thus, log2FC > 0 indicates higher abundance under Fe deficiency and log2FC < 0 indicates lower abundance. Unsupervised multivariate analysis by Principal Component Analysis (PCA) was conducted on the normalized (log10-transformed, mean-centered) data to visualize global differences in metabolite profiles between groups.

For Fe mobilization rate from ferric oxide and ferrihydrite, (n = 5), data were expressed as µg Fe·g⁻¹ ferric oxide·h⁻¹ and analyzed by one-way ANOVA followed by Tukey’s post hoc test to determine pairwise differences among groups. Statistical significance was accepted at p < 0.05. For Fe solubilization rate from ferric oxide in the presence of phenolic metabolite (n = 3), Fe concentrations were first normalized within each independent experiment to the median fraxetin value, which served as an internal positive control and was set to 1. Differences between individual treatments and the blank were evaluated using Welch’s t-tests, chosen to account for potential heterogeneity of variances; no correction for multiple comparisons was applied.

## 3 RESULTS AND DISCUSSION

### 3.1 Poplar responses to iron deficiency at alkaline pH

Two hydroponics experiments were carried out under controlled culture conditions, exposing poplars to Fe deficiency during 20 and 27 days. In order to monitor the progress of Fe deficiency, we measured the pH of the NS daily, before renewing it. Every 2-3 days the chlorophyll index (SPAD) was measured in the three upper leaves of the three plants of every pot of both treatments (control and Fe deficiency). Fig. 1 shows the pH and SPAD behavior during Fe deficiency. Data from the two experiments were pooled to assess trends.

**Fig. 1.**
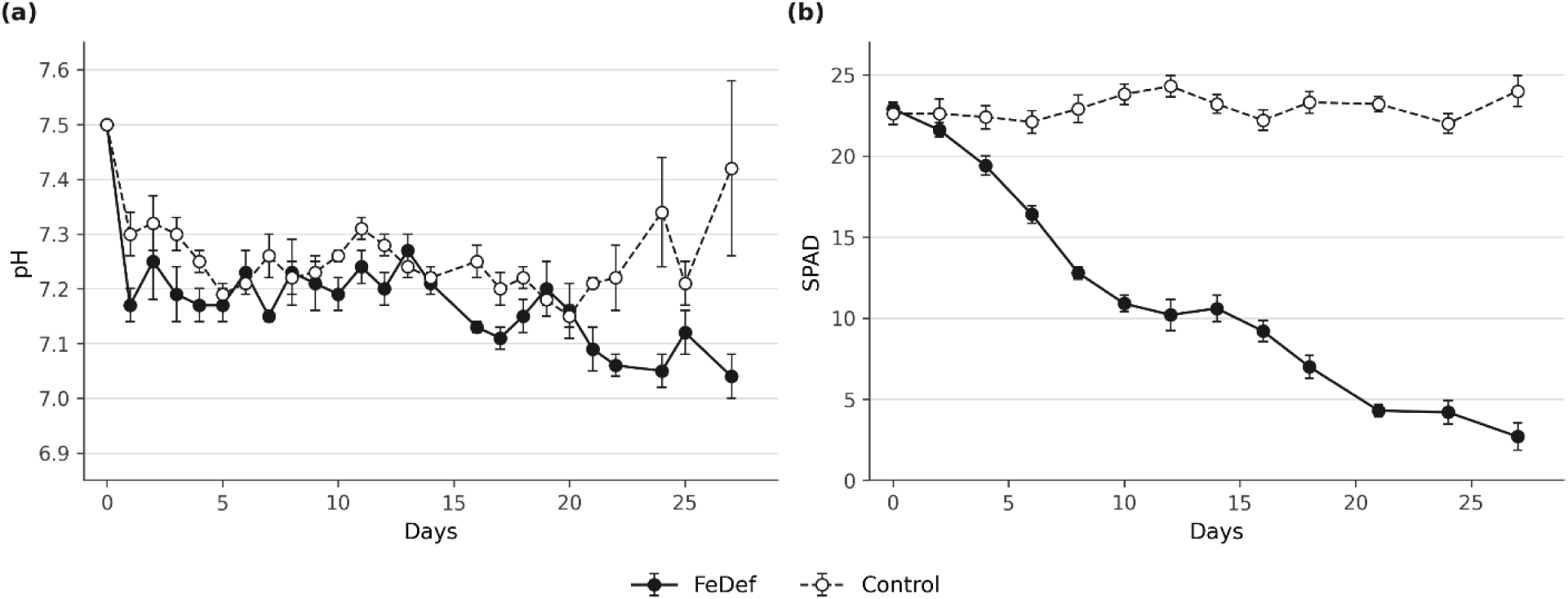
Time-course changes in nutrient solution pH (a) and leaf SPAD index (b) in Populus × canescens under Fe deficiency (FeDef) and control conditions over 27 days. pH was measured directly in the nutrient solution (five independent replicates per treatment). SPAD values were recorded on the three uppermost, fully expanded leaves of three plants per pot, with five pots per treatment used as independent biological replicates. Data are means ± standard error (n = 5) **Alt text:** Two line graphs over 27 days comparing Fe-deficient and control poplars. (a) Nutrient solution pH drops from 7.5 to about 7.2 in both treatments within the first day, then gradually declines to about 7.0 under Fe deficiency, while it stays between 7.2 and 7.4 in controls. (b) The leaf SPAD index stays at about 22–24 in controls but falls steadily under Fe deficiency, from about 23 to below 3 by day 27.

The pH of the deficient plants behaved similarly to the control plants up to 15 days and then tended to drop, with transient recoveries at 19 and 25 days. This is in agreement with the data reported by Masuda et al. (2018) showing that the poplar clone T89 decreases the pH of its growth medium upon Fe deficiency. We observed a continuous decrease of the SPAD index in Fe-deficient plants and a stable value in Fe-sufficient plants, as expected. This is consistent with the leaf chlorosis typically observed under Fe deficiency. Fig. 2 shows pictures illustrating the chlorosis developed by the plants after 20 and 27 days of Fe deficiency. Like other Strategy I plants, the leaf chlorosis observed in poplar is likely due to the inhibition of chlorophyll synthesis and photosynthesis. In fact, Chen et al. (2019) observed that, in Fe-deficient hybrid poplar clone 741, photosystem I was affected more severely and earlier than photosystem II.

**Fig. 2.**
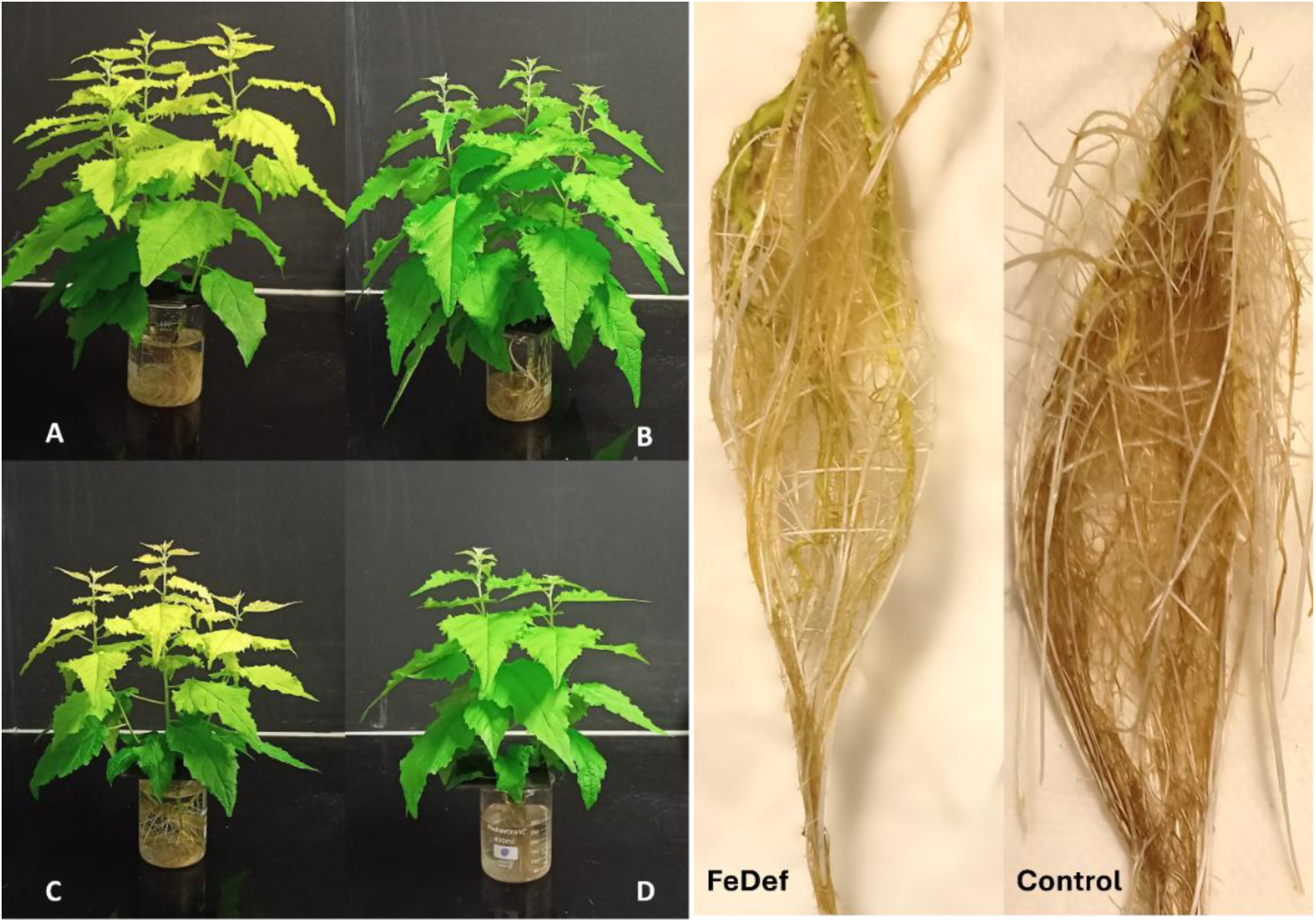
Iron deficiency symptoms. Left: pictures of Fe-deficient (A) and control plants (B) after 20 days and 27 (C and D) days of Fe deficiency. Right: picture of Fe-deficient and control roots after 27 days. **Alt text:** Photographs of hydroponically grown poplar plants and roots. Fe-deficient plants at 20 and 27 days have yellow (chlorotic) upper leaves, whereas control plants have uniformly green leaves. On the right, Fe-deficient roots look paler and more yellowish than the brown control roots after 27 days.

After 20 and 27 days under Fe deficiency, the plants were harvested and separated in leaves, stems and roots. The fresh weight of leaves, stems and roots and the length of stems are presented in Supplementary Table S2. We observed significant decreases between control and deficient leaves and stems fresh weights, indicating that Fe deficiency inhibited growth. In the case of roots, only the fresh weight from Fe deficient roots after 20 days presented a significant decrease with respect to control. Furthermore, no significant differences in stem height were observed between deficient and control plants, suggesting that, in response to a long-term Fe deficiency, poplars adopt the strategy to favor apical growth while decreasing stem, leaf and to a lesser extent root biomass. In contrast, Chen et al. (2019) observed that, in hybrid poplar clone 741, Fe deficiency promoted root growth, including increased root length and root number. The increased growth of the root system may be a strategy to improve Fe acquisition. *Populus tremula × Populus tremuloides* (T89) also showed increased root development and could survive long-term Fe deficiency. This genotype recovered rapidly upon Fe resupply, indicating robust Fe remobilization (Masuda et al., 2018). The alkaline pH used in our study, which probably led to more extreme Fe deficiency, may account for the discrepancy between our observations and those reported by Masuda et al. (2018) and Chen et al. (2019). No obvious differences were observed between control and deficient root architecture (Fig. 2-Right).

Fig. 3 shows a significant decrease in Fe concentration in leaves (left) and roots (right) of deficient compared to control plants at 20 and 27 days, as expected. Fe content in leaves of the deficient plants was 8.06 ± 0.17 mg kg^-1^ after 20 days and 5.78 ± 0.11 mg kg^-1^ after 27 days while in control plants was 18.5 ± 2.19 mg kg^-1^ after 20 days and 25.3 ± 2.25 mg kg^-1^ after 27 days. Fe concentration in roots of the deficient plants was 119 ± 18.5 mg kg^-1^ after 20 days and 91.7 ± 20.5 mg kg^-1^ after 27 days while in control plants was 448 ± 74.8 mg kg^-1^ after 20 days and 381 ± 77.4 mg kg^-1^ after 27 days.

**Fig. 3.**
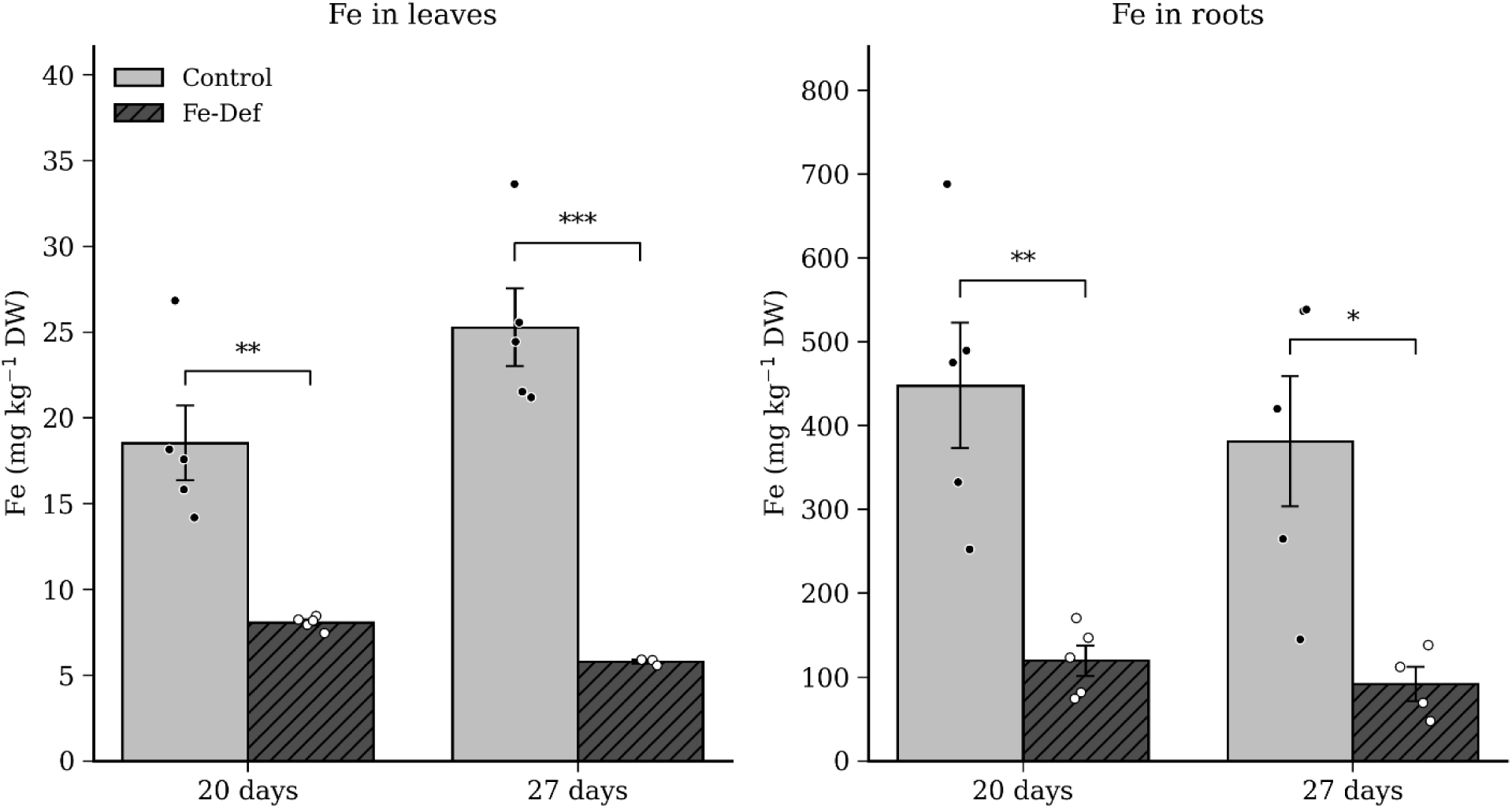
Iron concentration in leaves and roots of *Populus × canescens* at 20 and 27 days grown under control (Control) or iron deficiency (Fe-Def) conditions. Values are expressed as mg Fe kg⁻¹ dry weight (DW) and represent the mean ± standard error of the mean (SEM) of n = 3–5 biological replicates. Individual replicates are shown as dots (Control, filled; Fe-Def, open). Differences between treatments were evaluated by Student’s *t*-test after outlier detection by the iterative Grubbs test (α = 0.05), confirmation of normality (Shapiro-Wilk, *p* > 0.05) and homoscedasticity (Levene, *p* > 0.05). Asterisks denote statistically significant differences between Control and Fe-Def within each time point: *** *p* < 0.001, ** *p* < 0.01, * *p* < 0.05. **Alt text:** Two bar charts with individual data points showing Fe concentration in leaves (left) and roots (right) at 20 and 27 days. Fe-deficient plants have much lower Fe than controls at both times: about 8 and 6 versus 19 and 25 mg kg⁻¹ in leaves, and about 120 and 90 versus 450 and 380 mg kg⁻¹ in roots. All differences are significant.

To evaluate the behavior of other micronutrients, we measured zinc (Zn) and manganese (Mn) concentrations in leaves and roots of poplar after 20 and 27 days of Fe deficiency. The results presented in Supplementary Table S2 show no significant differences between control and deficient plants. In poplar, PtIRT1, which is mainly expressed in roots and strongly induced under Fe deficiency, particularly in Fe-chlorosis-resistant lines, has been proposed to drive Fe uptake at early stages of Fe deficiency (Huang and Dai, 2015). At later stages of Fe starvation, NRAMP1 becomes more strongly induced than IRT1, indicating a shift in metal uptake mechanisms (Chen et al., 2019). As IRT1 and NRAMP1 are not selective for Fe over other divalent metal cation, their induction enhances not only Fe uptake but can also promote the accumulation of other divalent metals, notably Zn and Mn, as reported in poplar and Arabidopsis. However, our results do not follow this trend. This could be due to the decreased Zn and Mn availability at alkaline pH.

As salicylic acid and jasmonic acid phytohormones are often affected by abiotic stress, we measured their concentrations in roots (Fig. 4). After 20 days of Fe deficiency, a non-significant decreasing trend in salicylic acid content in roots between control (2.09 ± 0.38 mg g^-1^) and deficient plants (1.17 ± 0.29 mg g^-1^) was observed (Fig. 4A). In contrast, after 27 days of Fe deficiency, there was a statistically significant increase in salicylic acid content in Fe deficient roots (1.98 ± 0.20 mg g^-1^) compared to control roots (0.99 ± 0.26 mg g^-1^, Fig. 4B). This indicates that poplar accumulates salicylic acid in roots upon long-term Fe deprivation. Jasmonic acid concentration was significantly higher in deficient roots (0.18 ± 0.02 mg g^-1^) than in control plants (0.07 ± 0.02 mg g^-1^) after 20 days of Fe deficiency (Fig. 4C). However, after 27 days of Fe deficiency, there was no longer any significant difference in jasmonic acid content between deficient (0.23 ± 0.11 mg g^-1^) and control roots (0.15 ± 0.03 mg g^-1^, Fig. 4D). These results suggest that jasmonic acid accumulates transiently in poplar roots when they are Fe deficient.

**Fig. 4.**
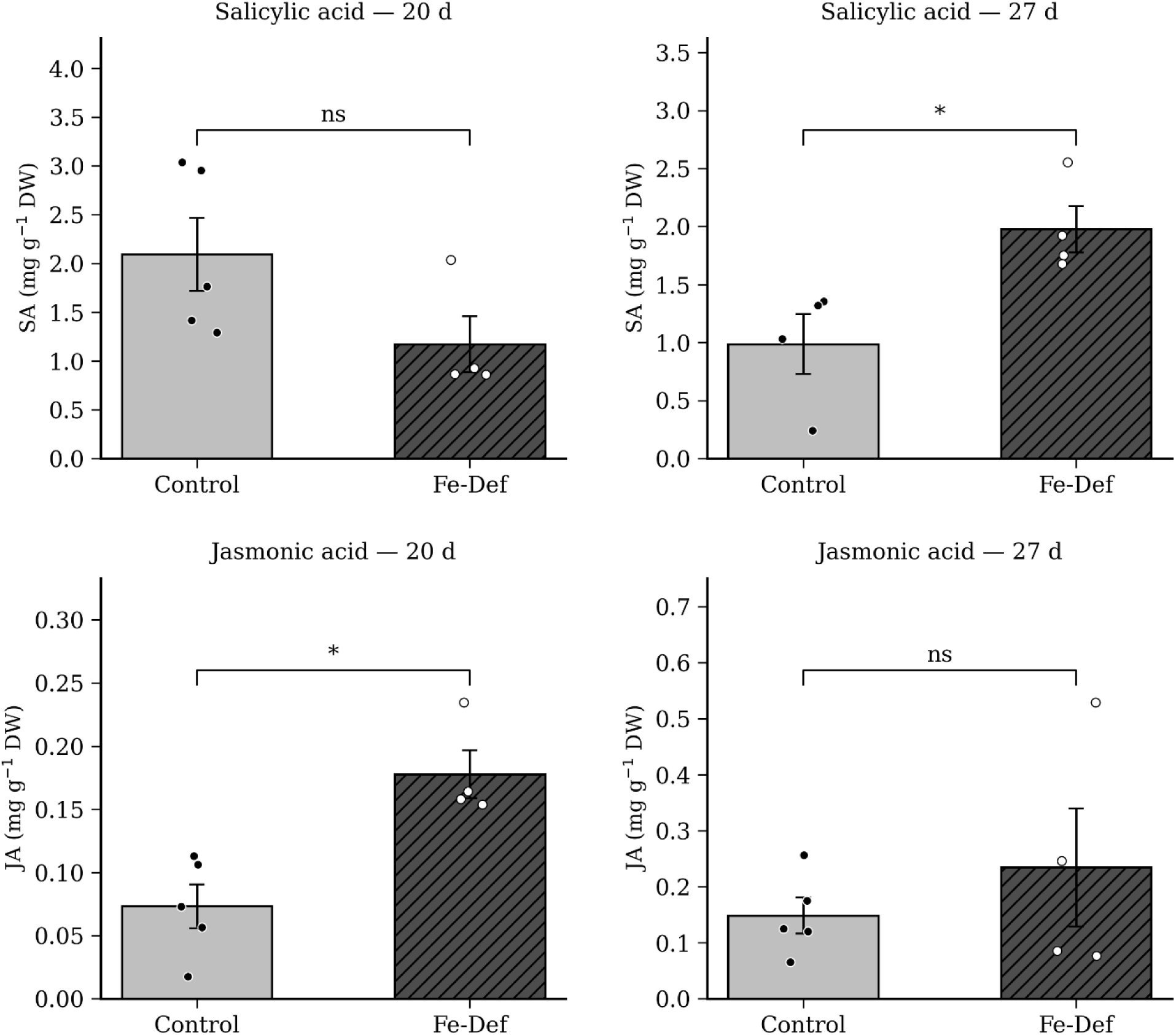
Salicylic acid (SA, upper panels) and jasmonic acid (JA, lower panels) concentrations in roots of *Populus × canescens* at 20 days (left) and 27 days (right) grown under control (Control) or Fe deficiency (Fe-Def) conditions. Values are expressed as mg hormone g⁻¹ dry weight (DW) and represent the mean ± standard error of the mean (SEM) of n = 4–5 biological replicates. Individual replicates are shown as dots (Control, filled; Fe-Def, open). Normality was assessed by the Shapiro-Wilk test and homoscedasticity by Levene’s test. Differences between treatments were evaluated by Student’s t-test when both assumptions were met, or by the Mann-Whitney U test otherwise. Asterisks denote statistically significant differences between Control and Fe-Def within each time point: * p < 0.05; ns, not significant. **Alt text:** Four bar charts with individual data points showing salicylic acid (top) and jasmonic acid (bottom) in roots at 20 and 27 days. Salicylic acid does not differ significantly at 20 days but is about twice as high in Fe-deficient roots at 27 days. Jasmonic acid is significantly higher in Fe-deficient roots at 20 days, with no significant difference at 27 days.

In contrast, auxins and abscisic acid concentrations in roots were not significantly affected by Fe deficiency either after 20 or after 27 days of starvation (Supplementary Table S2). As these hormones control root development, the finding that their levels are not changed is in agreement with the observation that Fe deficiency did not cause major alteration in root development. Jasmonic acid and salicylic acid are involved in stress signaling and may mediate some of the changes in the root metabolic profiles triggered by Fe deficiency (see below).

### 3.2 Effect of iron deficiency on root and exudate metabolomic profiles

In order to obtain as much information as possible about changes in the metabolic profile of poplar under Fe deficiency, we decided to collect root exudates in nutrient solution (RENS) and in distilled water (REDW). Exudates collected in a complete nutrient solution reflect a different metabolic state than in ion free water (Otxandorena-Ieregi et al., 2024). It is also likely that microorganisms contributed to decrease metabolite contents (Fracchia et al., 2024) since the experiments were carried out under non-sterile conditions. According to Oburger and Jones (2018), pure water (very low ionic strength) can rupture membranes, especially in young root hairs, releasing intracellular pools and leading to overrepresentation of “primary” metabolite families. Conversely, adding Ca²⁺ (common in nutrient solutions) stabilizes membranes but can stimulate re-uptake of amino acids, lowering is the amounts detected. In ionic media, transporters may actively take back certain metabolites (amino acids, sugars), changing the apparent profile; in water, steep gradients favor diffusion away from roots with less re-uptake. We tested the ability of RENS and REDW to mobilize Fe from insoluble sources, ferric oxide and ferrihydrite. RE extracts from Fe sufficient roots could not be tested due the high concentration of Fe in the extracts, which caused a high background. RENS increased Fe solubilization from ferric oxides by a factor 2 and 3 after 20 days and 27 days under Fe deficiency, respectively (Fig. 5). In contrast, REDW did not display any solubilization activity under these conditions. As, we suspected that the high background of solubilization from ferric oxides by the buffer alone could mask the activity of REDW, we tested whether REDW could solubilize Fe from a more stable source, ferrihydrite. Supplementary Fig. S1 shows that REDW extracts from poplars grown under Fe deficiency for 20 and 27 days increase Fe solubilization by 2 and 3 times compared to buffer, respectively. Unfortunately, due to limitation in the available material, we could not test RENS in these conditions. Collectively, our results show that RE extracts from Fe deficient poplars possess Fe solubilization activity, which varies according to the mode of collection and increases with the duration of the Fe starvation. This suggest that these extracts contain specialized metabolites with the potential to favor Fe solubilization from insoluble Fe sources.

**Fig. 5.**
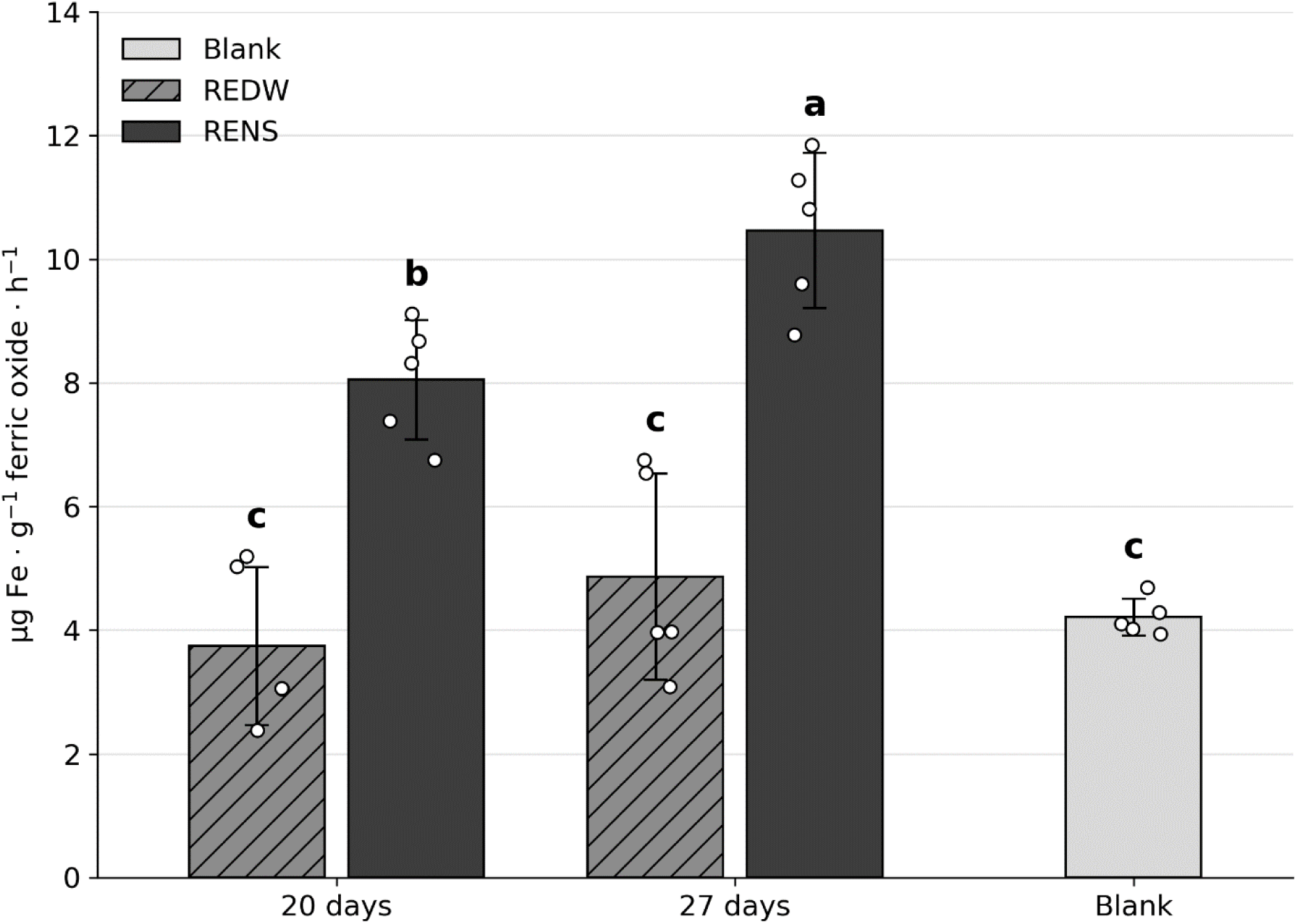
Iron mobilization rate from ferric oxide in the presence of root exudates collected in distilled water (REDW) and in nutrient solution (RENS). **Exudate**s were incubated for 6 h with 10 mg ferric oxide in HEPES buffer (pH 7.5) in the presence of NADPH and ferrozine. Iron concentrations were measured by MP-AES and expressed as µg Fe·g⁻¹ ferric oxide·h⁻¹. Values represent means ± SD (n = 5). The blank condition consisted of ferric oxide + NADPH + ferrozine without exudate. Individual replicates are shown as dots. Different letters indicate significant differences between groups (Tukey’s test, p < 0.05) **Alt text:** Bar chart of the Fe mobilization rate from ferric oxide by root exudates. Exudates collected in nutrient solution (RENS) mobilize more Fe than exudates collected in distilled water (REDW) and the blank, with the highest rate at 27 days. REDW does not differ from the blank at either time.

#### 3.2.1 Multivariate analysis of metabolomic profiles

The data matrix of molecular features (rt, *m/z*) for roots, root exudates in RENS and REDW samples were obtained after pre-processing the data and exported to the MetaboAnalyst 6.0 online software for multivariate analyses.

Principal component analysis (PCA), an unsupervised statistical multivariate method usually employed for exploratory data analysis and data visualization in untargeted metabolomics, was applied for each group of samples (Salem et al., 2022). Fig. 6 shows the results for roots, RENS and REDW. After 20 days of deficiency, the separation between the groups of deficient and control plants along PC1 was clear for roots, RENS and REDW, with more dispersion within each group along PC2. The total variance explained considering PC1 and PC2 for roots was 45.1%, for RENS 64.7 % and for REDW 79.1%. After 27 days of deficiency, we observed a clear separation along PC1 between treatments for RENS samples while, for roots and REDW, there is an obvious trend towards separation. The total variance at 27 days was 77.4% for RENS but only 38.5% for roots, and 39.2% for REDW. These results indicate clear separation between samples from control and Fe deficient plants albeit with a high biological heterogeneity, which is expected after 2 months of culture and quite common in untargeted metabolomic studies (Jeppesen and Powers, 2023).

**Fig. 6.**
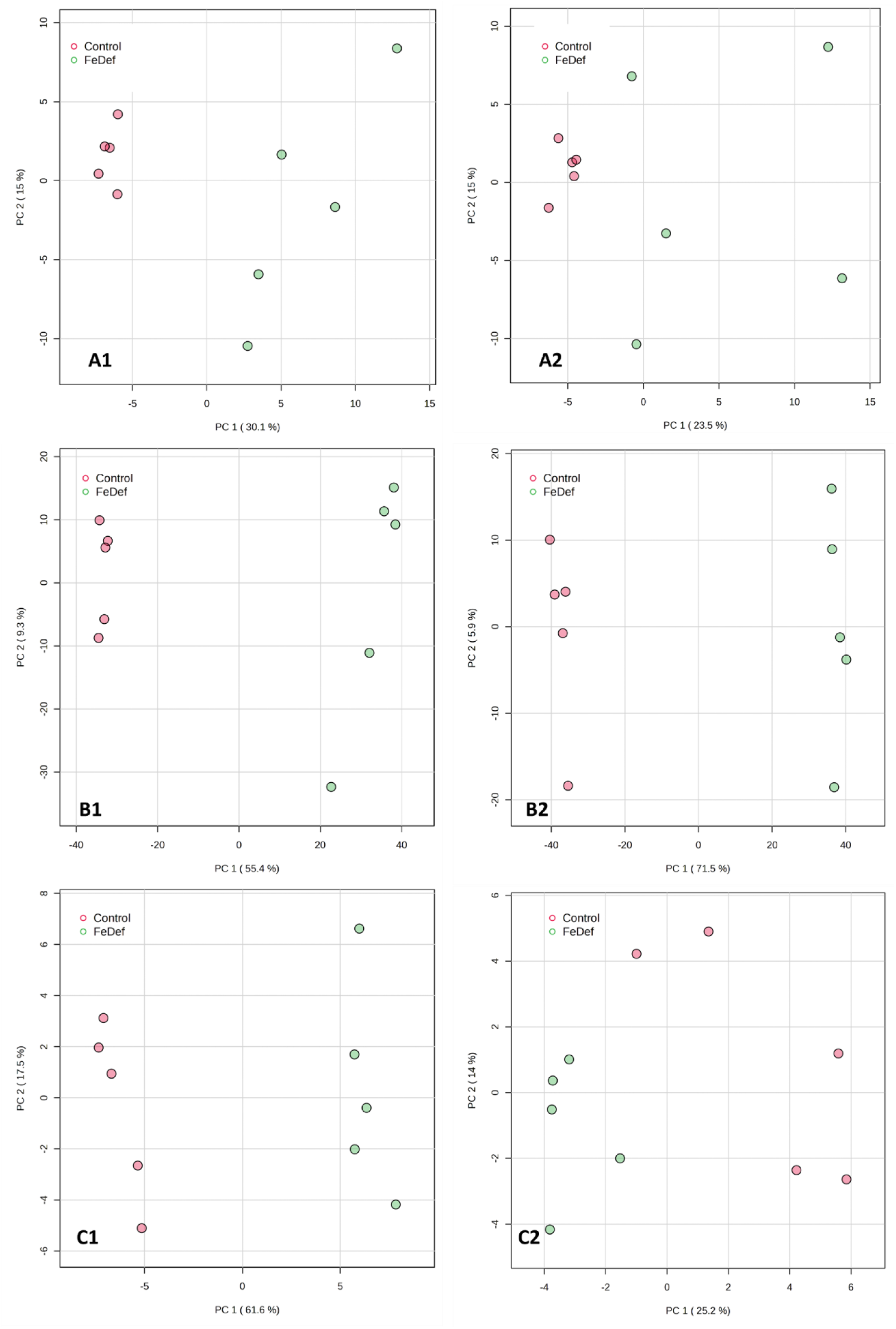
PCA analysis of molecular feature data from roots (A), root exudates in NS (B) and root exudates in DW (C) after 20 (1) and 27 days (2) of Fe deficiency. **Alt text:** Six principal component analysis score plots of metabolomic features from roots (A), exudates in nutrient solution (B) and exudates in distilled water (C) at 20 days (1) and 27 days (2). In most panels, samples from Fe-deficient and control plants form separate groups along the first principal component.

#### 3.2.2 Univariate analysis

To clarify the differences in metabolomic profiles between Fe-deficient and control plants, molecular features from roots, RENS and REDW were sorted according to t-tests (p < 0.05) and their log 2FC analysis. The log2FC values higher than 0 were considered increased and lower than 0, decreased. Table 1 summarizes the number of statistically significant and non-significant total and annotated features in the samples of roots, RENS and REDW after 20 and 27 days of Fe deficiency; the non-significant annotated metabolites are listed in Supplementary Tables S3–S8. We used the following criterion for metabolite annotation: the metabolite score must be equal to or higher than 0.7, which implies more than 70% agreement with the database integrated in MetGem (GNPS, MS-Dial, ISDB). However, we also annotated some compounds with lower database matches but which clustered with high-match metabolites in the molecular networks (Supplementary Figs S2–S7), assuming that these molecules belonged to the same family (clusters in Supplementary Figs S8–S10).

**Table 1:** Summary of statistically significant and non-significant total and annotated features in poplar samples of roots, RENS and REDW after 20 and 27 days of Fe deficiency. ND: not detected

|  |  | 20 days Fe deficiency |  | 27 days Fe deficiency |  |
| --- | --- | --- | --- | --- | --- |
|  |  | Features | Annotated | Features | Annotated |
| Roots | Total | 353 |  | 477 |  |
|  | Significant | 39 | 1 | 43 | 7 |
|  | Non-significant | 314 | 39 | 434 | 59 |
| RENS | Total | 649 |  | 513 |  |
|  | Significant | 318 | 3 | 291 | 8 |
|  | Non-significant | 331 | 22 | 222 | 9 |
| REDW | Total | 188 |  | 167 |  |
|  | Significant | 37 | ND | 65 | 1 |
|  | Non-significant | 151 | 7 | 105 | 7 |

According to Table 1, the number of features increased by around 35% in seven days. However, for the exudates in both nutrient solution and water, we observed the opposite trend, 21% less for RENS and 11% less for REDW. Root exudation profiles are dynamic and change over time, even within short periods like 4 to 30 days after planting (Fracchia et al., 2024). Moreover, our experiments were performed under non-sterile conditions, so it is possible that the decrease in the number of features in the nutrient solution could potentially result from microbial degradation as well as other abiotic processes occurring in the culture medium.

As can be noted in Table 1, between 11 and 14% of the features detected in root and around 4% of those detected in root exudates could be annotated. According to Gauglitz et al. (2022) only 10% of features in untargeted studies are confidently annotated, leaving most as unknowns. Although there has been great progress, spectral and structure databases cover only a fraction of possible metabolites, limiting confident matches (Chaleckis et al., 2019).

### 3.3 Metabolites in Roots

#### 3.3.1 Lipids

Table 2 shows metabolites with statistically different abundance between control and Fe-deficient plants in roots, RENS and REDW after 20 and 27 days of Fe deficiency. After 20 days, glyceryl linolenate increased significantly in roots, while at 27 days the difference was not significant (Supplementary Table S4). Glyceryl linolenate is a glycerolipid functionally related to alpha-linolenic acid that is a jasmonic acid precursor (Rehman et al., 2023) and its increase is coincident with the hormone increase in roots of Fe-deficient plants at 20 days (Fig. 4C). Jasmonic acid functions as a signaling molecule that modulates the expression of stress-responsive genes in response to biotic and abiotic stress conditions, for example, heavy metals toxicity. Notably, jasmonic acid promotes lignin deposition and secondary cell wall thickening in poplar trees through JAZ5-MYB/NAC interaction (Zhao et al., 2023).

**Table 2:**
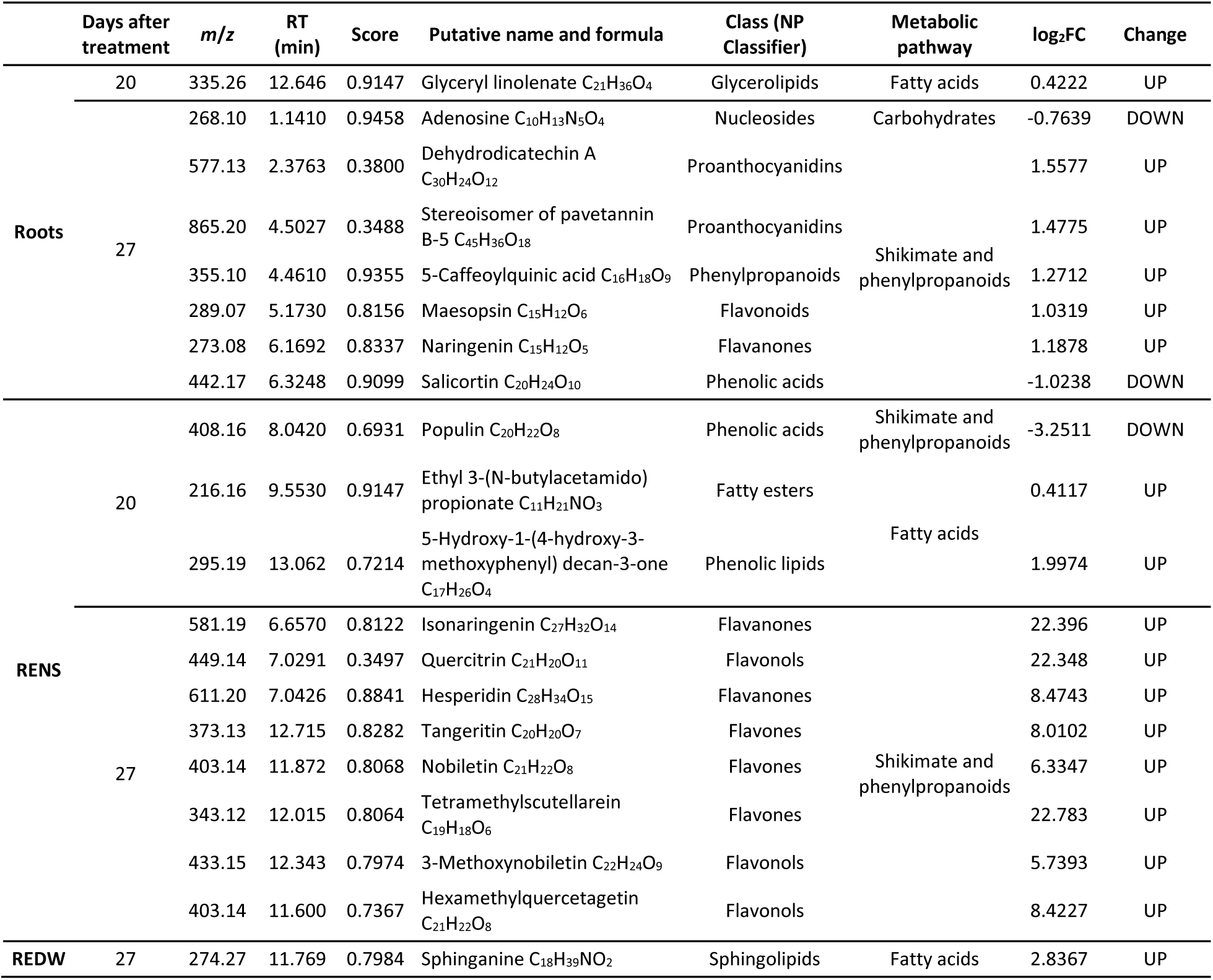
Significant metabolites annotated from roots, root exudates in nutrient solution (RENS) and root exudates in distilled water (REDW) after 20 and 27 days of Fe deficiency (p value<0.05). Change according to log2FC analysis (up means increase: log2FC>0, and down means decrease log2FC<0).

Sphinganine, a C18-sphingolipid (d18:0), was detected in root extracts in agreement with the measurement of increased levels of this metabolite in exudates (REDW) after 27 days.

#### 3.3.2 Phenylpropanoids

The phenylpropanoid pathway (Supplementary Fig. S11) is a central metabolic route that enables plants to adapt to biotic & abiotic stress by producing a diverse array of specialized metabolites, including flavonoids, phenolic acids, and proanthocyanidins (PAs), that play protective roles. After 27 days, PAs, flavonoids and flavanones significantly accumulated in the roots of Fe-deficient plants (Table 2). Interestingly, accumulation by 2-3-fold of dehydrodicatechin A, maesopsin and 5-caffeylquinic acid, also called neochlorogenic acid could already be detected after 20 days of Fe deprivation but they were not statistically significantly increased at this stage (Supplementary Table S3). Therefore, there was an increased accumulation of these compounds between 20 and 27 days of deficiency.

The biosynthesis of neochlorogenic acid diverges from the phenylpropanoid pathway at the caffeoyl-CoA node, which also feeds into lignin formation through the cinnamoyl-CoA reductase–cinnamyl alcohol dehydrogenase branch. Thus, neochlorogenic acid is not part of the lignin pathway per se, but it shares common precursors and competes for metabolic flux within the same network (Yao et al., 2021). Neochlorogenic acid plays a role in the defense mechanisms (Soviguidi et al., 2022) and stress responses of *Populus* species, such as *Populus tremula* under elevated temperature, enhanced UVB and UVA radiation (Nissinen et al., 2017).

The PAs are widespread polymeric flavan-3-ols known for their ability to bind proteins. They are synthesized via the flavonoid branch of the phenylpropanoid pathway, originating from phenylalanine (Supplementary Fig. S11). This biosynthetic pathway involves several enzymes including chalcone synthase, dihydroflavonol reductase, leucoanthocyanidin reductase and anthocyanidin reductase. These enzymes lead to the formation of catechin (2,3-trans) and epicatechin (2,3-cis), the monomeric building blocks of PAs (Dixon and Sarnala, 2020). The presence of catechin and epicatechin in roots was detected at both 20 and 27 days in both Fe sufficient and deficient conditions (Supplementary Tables S3, S4). Gourlay et al. (2022) found that in *Populus tremula × P. tremuloides* PAs production increases in leaves under oxidative stress from drought and UV-B. PAs are powerful antioxidants, protecting cellular components from oxidative damage caused by reactive oxygen species under abiotic stress. Then, increased levels of these antioxidants in poplar roots suggest a preventive reaction to oxidation and cell damage in response to Fe deficiency.

When the biosynthesis of monolignols is limited due to nutrient stress or genetic alterations, plants exhibit a metabolic adaptation in the phenylpropanoid pathway. Specifically, the carbon flux is redirected toward flavonoid metabolism, leading to the enhanced synthesis and accumulation of PAs, which can act as functional substitutes for lignin in cell wall reinforcement and antioxidant defense. These metabolites can polymerize and integrate into the cell wall, mimicking lignin in both structure and function, as shown in other contexts of phenylpropanoid flux reallocation (Wang et al., 2019; Zheng et al., 2024). Silencing of the lignin biosynthetic gene HCT (hydroxycinnamoyl-CoA:shikimate hydroxycinnamoyl transferase) in Arabidopsis thaliana or knocking out C3’H3 in Populus tomentosa redirects metabolic flux toward flavonoid biosynthesis, confirming the competitive regulation between lignin and flavonoid pathways (Besseau et al., 2007; Zhang et al., 2023). Transcriptional regulators such as MYB6, MYB165, and MYB194 in poplar have been shown to coordinate this balance (Ma et al., 2018; Wang et al., 2019). Transgenic tobacco plants experiencing Fe deficiency exhibit an decrease of lignin-related genes and increased accumulation of total flavonoids and proanthocyanidins (Kumar et al., 2013). In contrast, in *Populus ussuriensis*, the transcription factor PuC3H35 was shown to co-regulate both lignin and PA biosynthetic genes, promoting the accumulation of PAs for antioxidant protection and lignin for mechanical reinforcement under drought stress. This dual change indicates that the PAs and lignin pathways do not always compete but can be coordinately activated as part of an integrated stress response (D. Li et al., 2022), to Fe deficiency. Accordingly, Tadesse et al. (2025) showed that Fe deficiency leads to increased secondary cell wall deposition in *Populus x canescens* shoots.

After 27 days of Fe deficiency, the salicylic glycoside (SGs) salicortin (Supplementary Table S4) concentration decreased significantly by about 50% (Table 2). This is in agreement with the finding that, in poplar, increase in PAs biosynthesis correlates with decrease in SGs, (Liu et al., 2023). This is due to competition for shared precursors like cinnamic acid, which contributes to both salicyl and HCH (6-hydroxy-2-cyclohexan-1-carboxylate) moieties in salicortin (Babst et al., 2010).

After 27 days naringenin was significantly increased by 2-3-fold (Table 2). It is a crucial intermediate in the early steps of flavonoid biosynthesis (Supplementary Fig. S11). It is derived from chalcone via the enzyme chalcone isomerase, and serves as the immediate precursor for multiple downstream products including: dihydroflavonols, which feed into anthocyanins and PAs, flavonols, and flavanones (Yu et al., 2020). Interestingly, increased accumulation of a suite of flavonoids was detected in the RENS at the same time point (Table 2).

After 20 days, metabolites with well documented roles in Fe mobilization, fraxetin, fraxidin and, 5.8-dihydroxy-6-methoxy-7-[(2S.3R.4S.5S.6R)-3.4.5-trihydroxy-6-(hydroxymethyl) oxan-2-yl] oxychromen-2-one (sideretin glucoside – Robe et al., 2025) were detected in roots but no significant difference was observed between Fe sufficient and Fe deficient roots (Supplementary Table S3). After 27 days, in addition to the coumarins mentioned above, 5-OH-isofraxidin 7-O-glucoside, isofraxoside and skimmin were also detected (Supplementary Table S4). Moreover, all these coumarins except skimmin tended to accumulate in Fe-deficient roots at 27 days (log₂FC 1.0–2.1, i.e. about 2-to 4-fold; Supplementary Table S4), but the differences were not significant owing to high variability among biological replicates.

### 3.4 Metabolites from root exudates

#### 3.4.1 Lipids

Sphinganine was significantly increased by about 7-fold in REDW after 27 days of Fe deficiency (Table 2) confirming a trend already apparent after 20 days of Fe deficiency, more than 3% (Supplementary Table S7). Sphingolipids are a notable component of root exudates, and they play significant roles in plant response to various abiotic stresses. Fracchia et al. (2024) detected sphinganine, sphingosine and phytosphingosine in poplar exudates. In addition, Zhang et al. (2021) showed by non-targeted metabolomics (UHPLC/Q-TOF-MS) that the sphinganine content in root exudates of *Pennisetum americanum* exposed to atrazine, increased ∼5-fold after six days of exposure. The authors propose that the secretion of this long chain base (LCB) is part of the antioxidant response that helps protect the root against herbicide-induced damage by reactive oxygen species. However, the direct functional link between exuded sphingolipids and specific abiotic stress responses in the rhizosphere is not established yet. According to Panikashvili and Aharoni (2008), sphingolipid turnover releases LCBs and elevated levels of LCBs activate cell death pathways in *Arabidopsis*. The authors propose that Arabidopsis root cells export LCBs to maintain subtoxic concentrations, a strategy analogous to the export of cuticular waxes and other lipids via ABC transporters. Poplar may also exude LCBs to modulate the microbiota (Glenz et al., 2022), favoring stress tolerance-promoting microorganisms, as well as to signal systemic defense pathways (e.g. complexes with Ca²⁺/reactive oxygen species) (Seo et al., 2021). Then, the Fe-deficient poplar might exude sphinganine in order to avoid toxic LCBs accumulation in its roots, as well as to modulate microbiota and activate defense signals.

#### 3.4.2 Phenylpropanoids

After 20 days of Fe deficiency, populin amount dropped significantly to 10% of the value in Fe sufficient condition in RENS. M. Li et al. (2022) and Fracchia et al. (2024) also detected SGs in poplar root exudates, where they play significant roles in plant-microbe interactions and defense. The lower concentration of populin in exudates correlates with salicylate decrease in roots to promote PAs biosynthesis.

After 27 days of Fe deficiency, eight flavonoids (flavonols, flavones and flavanones) were highly significantly abundant in RENS by over 50-fold (Table 2). They already highly increased after 20 days but were not significantly different between Fe-deficient and Fe-sufficient exudates at this stage due to high variability among biological replicates (Supplementary Table S5). Flavonoids in root exudates may play several key protective and signaling roles. They act as crucial signaling molecules that modulate the interaction between roots and soil microbes as for example, beneficial plant-microbe symbioses (Yu et al., 2021). High concentrations of phenylpropanoids, including flavonoids, in roots or root exudates can be induced by nutritional stresses. Although flavonols are a major class of flavonoids found in *Populus* species (Chen et al., 2009; Zhong et al., 2012; Yang et al., 2023; Guleria et al., 2024), they had not yet been detected in root exudates. Specific flavonoids may bind and solubilize Fe in the rhizosphere: flavonols have a 3-hydroxyl (3-OH) group on the C-ring, which combines with the 4-oxo group to form a strong bidentate Fe-binding site (Butun et al., 2018). This feature is absent in flavones and flavanones, reducing their metal-binding capacity. Flavonols like quercetin and myricetin were found to strongly chelate Fe³⁺, while flavanones and flavanols were mostly inactive (Mira et al., 2002). Accordingly, among all the flavonoids listed in Table 2, quercitrin, putatively annotated on the basis of its clustering with hesperidin and isonaringenin in the molecular network (Supplementary Fig. S10), is the only one shown to act as a chelating agent (Li et al., 2016). Compared to its aglycone quercetin, the sugar moiety reduces but does not abolish chelation (Hu et al., 2006). Although 3-methoxynobiletin and hexamethylquercetagetin are also flavonols increased in exudates from Fe-deficient plants, their hydroxide groups are fully methylated, which abolishes Fe chelation (Lomozová et al., 2021). Based on this knowledge, we tested the possibility that quercitrin could solubilize Fe from insoluble sources and could therefore account, at least in part, for the Fe solubilizing activity of RENS (Fig. 5). To this aim, we compared the Fe solubilizing activity of quercitrin with quercetin, its non-glycosylated moiety, fraxetin, a well characterized Fe solubilizing metabolite of Fe deficient exudates, and the flavanone hesperidin (Fig. 7). As expected, hesperidin did not solubilize more Fe from ferric oxides than the control buffer. In contrast, the flavonol aglycone quercetin allowed increased solubilization by a factor 5-6. Quercitrin displayed an intermediate activity, comparable to that of the well-characterized fraxetin, increasing Fe solubilization by a factor 2-3. Qualitatively similar results were obtained when using ferrihydrite instead of ferric oxides as non-available Fe substrate (Supplementary Fig. S12). Together, these results identify quercitrin as a metabolite that highly accumulates in poplar root exudates under Fe deficiency and is capable of increasing Fe availability in the rhizosphere.

**Fig. 7.**
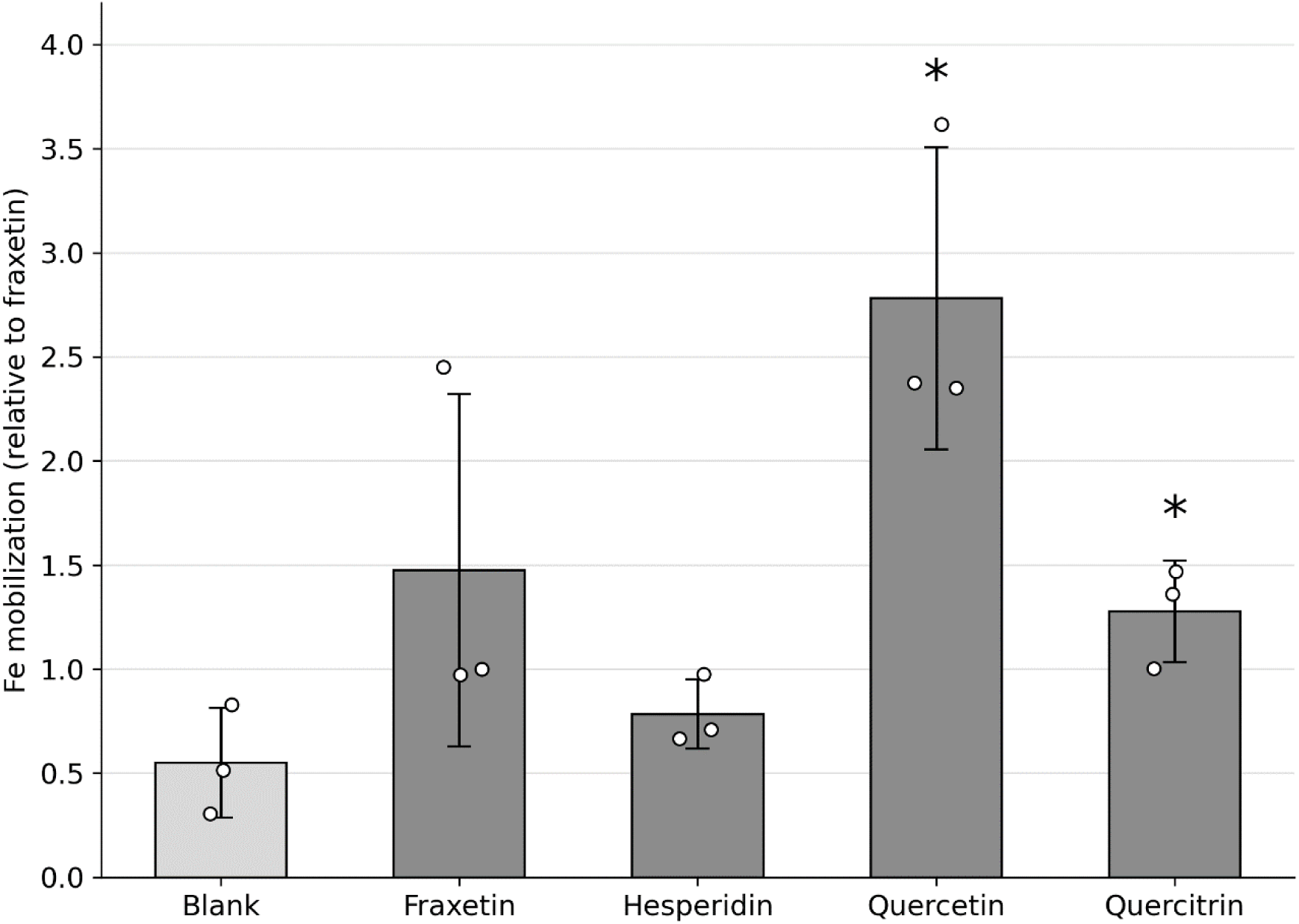
Iron solubilization rate from ferric oxide in the presence of phenolic metabolites. Extracts were incubated for 6 h with 10 mg of ferric oxide in HEPES buffer (pH 7.5) in the presence of NADPH and ferrozine. Iron concentrations were quantified by MP-AES and values were normalized within each experimental set to the median fraxetin value (internal positive control), which was set to 1. Data represent mean ± SD (n = 3). The blank condition consisted of ferric oxide + NADPH + ferrozine without metabolite. Individual replicates are shown as dots. Statistical comparisons were performed using Welch’s t-tests without correction for multiple comparisons. **Alt text:** Bar chart of Fe mobilization from ferric oxide by individual phenolic compounds, expressed relative to fraxetin. Quercetin gives the highest mobilization (about 2.8) and quercitrin about 1.3; both are significantly higher than the blank. Fraxetin and hesperidin do not differ significantly from the blank.

In exudates, coumarins were detected only in REDW. No clear pattern of coumarin exudation appeared after 20 days of Fe deficiency, with non-significant increases or decreases (e.g. fraxetin, log₂FC −3.1). After 27 days, fraxetin and isofraxidin were about 10-fold more abundant in REDW from Fe-deficient plants (log₂FC 3.6 and 3.4, respectively), and 6,7-dimethoxycoumarin was detected almost exclusively in exudates of Fe-deficient plants (log₂FC 19.5) (Supplementary Tables S7, S8). However, these differences did not reach statistical significance, because of high variability among biological replicates. The identity of the main coumarins was confirmed by a complementary LC–MS analysis of pooled root and REDW samples (Supplementary Table S9). Fraxetin, fraxin, isofraxidin and fraxinol (6-hydroxy-5,7-dimethoxycoumarin) matched authentic standards in retention time, accurate mass and MS/MS spectrum. Fraxetin and fraxinol were detected in both roots and REDW, isofraxidin in roots and, only under Fe deficiency, in REDW, and fraxin only in roots. All these coumarins gave higher signals in samples from Fe-deficient plants than in controls (1.5-to 2.3-fold in roots and 1.7-to 2.1-fold in REDW). In contrast, scopoletin, the precursor of fraxetin, was not detected in any sample, which may reflect its efficient conversion into catecholic coumarins in poplar roots. Because this analysis was performed on pooled samples, it confirms the identity of these coumarins but does not allow statistical comparison between treatments; nevertheless, it is consistent with the trend towards higher coumarin levels under Fe deficiency observed in the untargeted analysis.

In *Arabidopsis thaliana*, the function of coumarins in Fe acquisition is highly dependent on environmental pH. In alkaline conditions (pH 7.5), the predominant metabolite shifts from sideretin to fraxetin, which primarily mobilizes soluble Fe (III) for subsequent reduction by FRO2, rather than directly reducing Fe (III) itself (Paffrath et al., 2024). Poplar genomes encode genes homologous to those encoding the key enzymes involved in coumarin biosynthesis and the ABCG37/PDR9 transporter involved in coumarin secretion in Arabidopsis (Masuda et al., 2018). Although Masuda et al. (2018) did not explicitly identify coumarins in poplar root exudates, they showed that coumarin biosynthesis and export genes are upregulated under Fe deficiency along with genes encoding the Fe uptake machinery (e.g., *PtIRT1*, *PtFRO2*). Using a multi-omics approach, Zhang et al. (2024) showed that coumarin accumulation in *Populus tomentosa* leaves, including fraxetin, is genetically controlled and shows signatures of local adaptation, mainly associated with climatic variables such as temperature and precipitation. Liu et al. (2024) detected fraxetin in the rhizosphere soil of field-grown poplar hybrids, with higher levels in some offspring genotypes than in their parents. These studies indicate that coumarin accumulation varies among poplar genotypes; together with our results and with studies in Arabidopsis and tobacco, it is likely that poplars produce coumarins in their roots and exude them into the rhizosphere as a strategy to counteract Fe deficiency. However, our results failed to demonstrate a statistically significant increase in coumarin production and exudation, under Fe deficiency in contrast with flavonoid production.

Flavonoids and coumarins are produced in response to Fe deficiency via separate branches of the phenylpropanoid pathway (Robe et al., 2021; Rosenkranz et al., 2021). It has been widely demonstrated that coumarins are the primary Fe-mobilizing and chelating agents in the rhizosphere of Arabidopsis, especially under Fe-limiting and alkaline soil conditions (Fourcroy et al., 2014; Tsai and Schmidt, 2017; Robe et al., 2021; Rosenkranz et al., 2021; Paffrath et al., 2024; Robe et al., 2025). Flavonoids, while chemically capable of Fe chelation and reduction, have so far mostly been considered as signaling molecules for plant-microbe interactions and as modulators of the rhizosphere microbiome (Cesco et al., 2010, 2012; Kejík et al., 2021; Kumar et al., 2024). However, Terzano et al. (2015) showed that flavonoids are not merely signaling molecules but can act as potent Fe-mobilizing agents in calcareous soils, especially when combined with organic acids. Our results demonstrating that, among the flavonoids increased under Fe deficiency in poplar exudates, quercitrin displays Fe solubilization capabilities, support this conclusion.

## 4 CONCLUSIONS

Iron deficiency under alkaline conditions induces a global metabolic reprogramming in *Populus × canescens*, reshaping both root and root exudates metabolomes over a 20–27-day time course. Beyond classical physiological responses, Fe deprivation redirects key secondary metabolic pathways, particularly the phenylpropanoid–flavonoid–proanthocyanidin axis, and modifies rhizosphere composition through the selective release of specialized metabolites.

Fe deprivation progressively reprogrammed the root metabolome, redirecting the phenylpropanoid pathway toward proanthocyanidins, neochlorogenic acid, and naringenin at the expense of salicylate glycosides—consistent with a metabolic trade-off between lignin/salicylate and flavonoid branches. This shift was likely hormonally coordinated, with transient jasmonic acid accumulation at 20 days and significant salicylic acid increase at 27 days. Catecholic coumarins (fraxetin, fraxin, isofraxidin and fraxinol) were detected in roots and root exudates, and their identity was confirmed with authentic standards; although they tended to accumulate under Fe deficiency, this increase could not be validated statistically. In contrast, flavonoids were highly and robustly increased in nutrient-solution exudates after 27 days.

By integrating untargeted UHPLC–QTOF metabolomics coupled with MS/MS molecular networking, confirmation with authentic standards and functional Fe-solubilization assays, this study demonstrates that poplar root exudates possess the capacity to mobilize Fe from poorly available sources at high pH. While catecholic coumarins are established contributors to Fe acquisition, our results indicate that specific flavonoids—such as quercitrin—can also actively participate in Fe solubilization, expanding the current framework beyond coumarin-centered models and highlighting a more diverse chemical strategy.

Only about 10% of detected metabolite features could be confidently annotated, underscoring both the current limitations of metabolite databases and the unexplored diversity of root-associated chemistry. Rather than weakening the conclusions, this highlights the potential of untargeted metabolomics in complex plant systems.

Overall, this work provides an integrated view of how alkaline Fe deficiency reconfigures root and exudate metabolomes in a model tree species and identifies root metabolites as active drivers of Fe mobilization in the rhizosphere.

These findings expand our understanding of Fe acquisition strategies beyond herbaceous model species and provide a metabolomic framework for future studies on tree crop adaptation to calcareous soils.

## Abbreviations

ABA: abscisic acid
DW: dry weight
ESI: electrospray ionization
FC: fold change
GNPS: Global Natural Products Social Molecular Networking
HBED: *N*,*N*′-bis(2-hydroxybenzyl)ethylenediamine-*N*,*N*′-diacetic acid
HEPES: 4-(2-hydroxyethyl)piperazine-1-ethanesulfonic acid
IAA: indole-3-acetic acid
ISDB: *in silico* spectral database
JA: jasmonic acid
LC-MS: liquid chromatography-mass spectrometry
MP-AES: microwave plasma atomic emission spectroscopy
MS/MS: tandem mass spectrometry
NADPH: reduced β-nicotinamide adenine dinucleotide phosphate
PA: proanthocyanidin
PCA: principal component analysis
PVDF: polyvinylidene fluoride
QTOF: quadrupole time-of-flight
REDW: root exudates collected in distilled water
RENS: root exudates collected in nutrient solution
RT: retention time
SA: salicylic acid
SEM: standard error of the mean
SG: salicylate glycoside
UHPLC: ultra-high-performance liquid chromatography

## Supplementary data

The following supplementary data are available at JXB online.

Table S1. Composition of the media used for in vitro propagation of poplar.

Table S2. Fresh weight of leaves, stems and roots, stem length, Zn and Mn concentrations, and auxin and abscisic acid contents in control and Fe-deficient plants after 20 and 27 days of Fe deficiency.

Table S3. Non-significant metabolites annotated in poplar roots after 20 days of Fe deficiency.

Table S4. Non-significant metabolites annotated in poplar roots after 27 days of Fe deficiency.

Table S5. Non-significant metabolites annotated in root exudates collected in nutrient solution after 20 days of Fe deficiency.

Table S6. Non-significant metabolites annotated in root exudates collected in nutrient solution after 27 days of Fe deficiency.

Table S7. Non-significant metabolites annotated in root exudates collected in distilled water after 20 days of Fe deficiency.

Table S8. Non-significant metabolites annotated in root exudates collected in distilled water after 27 days of Fe deficiency.

Table S9. Coumarins identified by complementary LC–MS analysis of pooled root and REDW samples from control and Fe-deficient plants.

Fig. S1. Iron mobilization rate from ferrihydrite in the presence of root exudates collected in distilled water (REDW).

Fig. S2. Molecular network of specialized metabolites annotated in poplar roots after 20 days of Fe deficiency.

Fig. S3. Molecular network of specialized metabolites annotated in poplar roots after 27 days of Fe deficiency.

Fig. S4. Molecular network of specialized metabolites annotated in root exudates in nutrient solution after 20 days of Fe deficiency.

Fig. S5. Molecular network of specialized metabolites annotated in root exudates in nutrient solution after 27 days of Fe deficiency.

Fig. S6. Molecular network of specialized metabolites annotated in root exudates in distilled water after 20 days of Fe deficiency.

Fig. S7. Molecular network of specialized metabolites annotated in root exudates in distilled water after 27 days of Fe deficiency.

Fig. S8. Cluster of proanthocyanidins in roots after 20 days of Fe deficiency.

Fig. S9. Cluster of proanthocyanidins in roots after 27 days of Fe deficiency.

Fig. S10. Cluster of flavonoids in root exudates collected in nutrient solution after 27 days of Fe deficiency.

Fig. S11. Phenylpropanoid pathway and its key enzymes.

Fig. S12. Iron mobilization rate from ferrihydrite in the presence of phenolic metabolites.

## ACKNOWLEDGEMENTS

This work has benefited from the support of IJPB’s Plant Observatory platforms P0-Chem. The IJPB benefits from the support of Saclay Plant Sciences-SPS (ANR-17-EUR-0007).

## AUTHOR CONTRIBUTIONS

MTC: conceptualization, methodology, investigation, formal analysis, data curation, writing – original draft, funding acquisition; NB: investigation (iron mobilization assays); SC: investigation and formal analysis (hormone analysis); DT: methodology, formal analysis (metabolomics), writing – review & editing; AAF: conceptualization, methodology; ST: conceptualization, supervision, writing – original draft, writing – review & editing, funding acquisition.

## DATA AVAILABILITY

The raw UHPLC–QTOF MS data (Agilent 6546) underlying the untargeted metabolomic analyses are deposited in Zenodo at https://doi.org/10.5281/zenodo.23265925 and will be made openly available upon publication. The complementary LC–MS data used to support metabolite annotation (Bruker compact QTOF) are available from the corresponding authors upon reasonable request. All other data supporting the findings of this study are available within the article and its supplementary data.

